# Quantifying local malignant adaptation in tissue-specific evolutionary trajectories by harnessing cancer’s repeatability at the genetic level

**DOI:** 10.1101/401059

**Authors:** N Tokutomi, C Moyret-Lalle, A Puisieux, S Sugano, P Martinez

## Abstract

Cancer is a potentially lethal disease, in which patients with nearly identical genetic backgrounds can develop a similar pathology through distinct combinations of genetic alterations. We aimed to reconstruct the evolutionary process underlying tumour initiation, using the combination of convergence and discrepancies observed across 2,742 cancer genomes from 9 tumour types. We developed a framework using the repeatability of cancer development to score the local malignant adaptation (LMA) of genetic clones, as their potential to malignantly progress and invade their environment of origin. Using this framework, we found that pre-malignant skin and colorectal lesions appeared specifically adapted to their local environment, yet insufficiently for full cancerous transformation. We found that metastatic clones were more adapted to the site of origin than to the invaded tissue, suggesting that genetics may be more important for local progression than for the invasion of distant organs. In addition, we used network analyses to investigate evolutionary properties at the system-level, highlighting that different dynamics of malignant progression can be modelled by such a framework in tumour-type-specific fashion. We find that occurrence-based methods can be used to specifically recapitulate the process of cancer initiation and progression, as well as to evaluate the adaptation of genetic clones to given environments. The repeatability observed in the evolution of most tumour types could therefore be harnessed to better predict the trajectories likely to be taken by tumours and pre-neoplastic lesions in the future.

## Introduction

Cancer is a disease fuelled by somatic evolution (Nowell, 1976), in which cells from multicellular organisms switch to uncurtailed growth (Davies & Lineweaver, 2011), potentially leading them to invade distant organs and eventually to death. This evolution is at least partly explained by the accrual of (epi)genetic alterations in cells over several generations and is context-specific, with different recurrent alterations observed in different tissue types (Greaves & Maley, 2012). The hypothesis that malignant transformation requires multiple alterations provides an explanation to why cancer incidence increases with age (Armitage & Doll, 1954) and why progression to cancer occurs via intermediate benign stages (Fearon & Vogelstein, 1990; Vogelstein et al., 2013). This is further corroborated by the observation that most solid adult tumours harbour multiple “driver” alterations, i.e. those likely to functionally impact cell behaviour and push it towards malignancy (Zack et al., 2013). Yet, the dynamics and stochastic nature of malignant transformation are still insufficiently understood: clinicians still lack efficient diagnostic tools to accurately predict if and when benign lesions will progress to cancer (Martinez et al., 2018), which patients at risk will develop tumours and what their genetic characteristics will be (Lässig, Mustonen, & Walczak, 2017).

The heterogeneity observed between tumours of the same type demonstrates that multiple evolutionary trajectories can converge toward malignancies with similar phenotypic characteristics (Yates & Campbell, 2012). Yet, how much impact individual driver alterations have on this adaptation or how they interact is still largely unknown. In addition, although metastatic cancer still represents a major clinical challenge, the role of the genetic makeup of the primary site in the adaptation to a novel distant site is also unclear. While reconstructing the genetic history of individual patients through sequencing becomes easier (Gerlinger et al., 2012; Nik-Zainal et al., 2012; Yates et al., 2017), the generic process of carcinogenesis underlying each tumour type remains elusive. The availability of public datasets describing the genetic characteristics of multiple specimen of various tumour types however represents an opportunity to decipher oncogenesis as a stochastic process. All tumours characterized so far are outcomes of their type-specific evolutionary process. This is akin to Gould’s contingency definition (Gould, 1990), as if “replaying the tape” of somatic evolution: recording all malignant developments starting from the mostly identical genetic backgrounds of different human individuals (Rosenberg et al., 2002).

Here we investigate these recorded outcomes to infer the evolutionary landscape of each tumour type, mapping the evolutionary trajectories that can lead normal cells to cancerous transformation. We make the following assumptions 1) cancer is of monoclonal origin, whereby the genetic background of the initial clone is detectable in all subsequent generations; 2) although different evolutionary trajectories can lead to such a clone, they all are similarly capable of invading their organ-specific environment of origin. We thus estimate different evolutionary parameters to investigate the contribution of all drivers within genetic clones and calculate a local malignant adaptation (LMA) score resulting from their combination in 9 tumour types. Similar to a fitness definition in evolutionary biology, our score can be understood as a measure of adaptation to a disease-specific evolutionary context, leading to harmful over-proliferation and domination of the local environment. Our model highlights differences across tumour types regarding the interactivity between driver alterations and predicts that pre-malignant skin and colorectal lesions are adapted to their environment, yet not as much as invasive tumours. We find that genetic landscapes of local adaptation do not explain the location of distant metastases, suggesting that adaptation to metastatic sites does not rely on genetics as much as tumorigenesis does. Finally, we suggest that networks can be used to represent stepwise genetic progression, providing useful tools to stochastically reconstruct the contingencies of the oncogenesis process and study its systemic properties.

## Methods

### TCGA data

We downloaded data for 2,742 samples from The Cancer Genome Atlas (TCGA), for which we could obtain both allelic frequencies for mutations and copy number alterations (CNAs). This represented 133 bladder cancers (BLCA), 914 breast cancers (BRCA), 195 colorectal cancers (COAD), 256 glioblastomas (GBM), 296 head & neck squamous cell cancers (HNSC), 306 kidney clear cell renal cell carcinomas (KIRC), 262 lung adenocarcinomas (LUAD), 132 lung squamous cell cancers (LUSC) and 248 skin melanomas (SKCM). Raw SNP array data were normalised using the aroma R package (Ortiz-Estevez, Aramburu, Bengtsson, Neuvial, & Rubio, 2012) with paired normals and copy number profiles were called using ASCAT (Van Loo et al., 2010).

### Nevi & Melanoma data

The mutational and copy number data for 37 primary melanomas with paired precursor lesions were downloaded from Shain et al (Shain et al., 2015). Nevi, blue nevi, and intermediate benign annotations were considered benign lesions, while intermediate malignant, melanoma in situ, melanoma (all stages), desmoplastic melanoma and blue-nevus-like melanoma annotations were considered malignant. In the few cases in which multiple malignant lesions were found for a patient, the least advanced was selected. Mutations were considered clonal when the normalised MAF values in the reported sequencing data was > 0.8. To detect CNAs, we first calculated the mean and standard deviation in segment mean (i.e. logR ratios for expected chromosomal copies in a segment) for all segments from the reported normal samples. Gains and losses for the 4 CN drivers retained for the melanoma landscape (CDK4_gain, CDKN2A_loss, MDM2_gain, PTEN_loss) were calculated based on the reported segment mean for the relevant segment of each sample deviated significantly from the normal segment mean distribution, using a one-tailed test with the pnorm R function. It is whoth noting that, unlike whole-exome TCGA data, these data are mostly obtained through targeted sequencing of a 293-gene panel and may therefore miss information on given drivers.

### Colorectal adenoma and carcinoma data

We retrieved whole-genome sequencing data from 31 samples from 9 adenomas and 72 samples from 11 carcinomas (*Cross et al. accepted at Nature Eco & Evo*). Mutations were called with platypus (Rimmer et al., 2014), copy number data and ploidy were obtained using the cloneHD software (Fischer, Vázquez-García, Illingworth, & Mustonen, 2014). Mutation clonality was estimated using the EstimateClonality package (McGranahan et al., 2015) on each of the 31 samples as for the TCGA data. Mutations were considered clonal in an entire adenoma when they were predicted as clonal in >= 75% of the related samples.

### MET500 data

Genomic data were retrieved from Robinson et al (Robinson et al., 2017) and via the MET500 website (https://met500.path.med.umich.edu/). Sample annotation was manually curated to attribute a tumour type to the primary site and metastatic site. All curated annotations are reported in Supp. Tables 2 and 3. Sample Purity was collected from the Supp. information of the original publication; sample ploidy was calculated as the mean copy number weighed by number of targeted exons. Mutation clonality was estimated using the EstimateClonality package.

### Clonality and driver alterations

Mutation clonality was estimated thanks to the EstimateClonality package (McGranahan et al., 2015), clonal mutations were defined as those for which the 95% confidence interval of the cancer cell fraction (CCF) included 1. Gain and losses of each gene in each sample were defined relatively, if the copy number deviated by 0.6 or more from the ploidy given by ASCAT in the segment that included the gene of interest. Segments with less than 10 probes were filtered out. Driver alterations for each tumour type were retrieved from the IntOGen website (Gonzalez-Perez et al., 2013; Rubio-Perez et al., 2015). Due to the lack of an appropriate clonality estimation method, all copy number alterations were considered clonal. Final matrices of clonal mutations per sample are available via github. To limit the number of potential combinations, only the 50 most frequent driver alterations that occured in > 0.5% of a cohort were selected. Due to their proximity on the chromosome, CDKN2A loss and CDKN2B loss were merged into a single event, which avoids later biases when investigating co-occurrence. This left 34 drivers in GBM, 47 in KIRC and 50 in all other tumour types.

### Quantification of evolutionary parallelism

We represented each datasets as a gene per patient matrix, using 0 to denote the absence of a clonal alteration and 1 for the presence of a specific clonal alteration in a specific patient. When multiple point mutations occur in the same gene in a patient, they are reported with a value of 1 encompassing all mutations. For each tumour type, 10 randomised matrices were generated by randomly reassigning the mutations of each patient. For instance, upon simulation, a patient with 300 clonal mutations out of 17,000 potential genes will still have 300 clonal mutations albeit in different genes than the originally observed alterations. The Jaccard Index between two samples is then computed using the following formula:

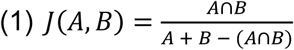

Where A and B are the number of alterations in each sample, and A∩B the number of overlapping alterations. All pairwise indices are computed for each matrix. Indices from the 10 simulated matrices are then pooled as randomised controls for a given tumour type.

### Genetic parameters of Local Malignant Adaptation

The selective advantage of each driver alteration was defined as the ratio between expectations and observations in a given tumour type, considering mutations and CNAs separately. The expected number of mutations occurring in a gene was calculated by dividing the total number of mutations by the total number of genes in which at least 1 mutation was observed, weighted by each gene’s length in base pairs, as given by the median transcript length for the gene in Ensembl (Zerbino et al., 2018). Only clonal mutations were considered. CNA selective advantage was calculated separately for gains and losses, by the ratio of expected occurrences given all events to the actual occurrences of each CNA, giving equal probability to all genes without weighing for length. The selective advantage *SAd* of each driver *d* is thus centred around 1 with a 0 lower boundary.

We hypothesised that drivers that tend to occur with few other drivers had more impact on malignant progression, and thus more “self-sufficient”, that those occurring with more additional drivers. To calculate self-sufficiency, the number of “partners” of each driver alteration was first computed. For each driver alteration *d* in a tumour type, this is given by the number of additional clonal drivers in *S*_*d*_, the subset of samples in which *d* is clonally mutated. For *d, Np*_*d*_ is thus a vector of number of partners of the same size as *Sd*. Defining *Np*_*nd*_ as the number of partners of all other drivers (“non *d*”) in all the samples in which they are mutated, we then compute *P*_*d*_, the power of finding such difference in mean between *Np*_*d*_ and *Np*_*nd*_ using the power.t.test R function, with *Np*_*nd*_ as the reference distribution. As self-sufficiency is inversely correlated to the number of partners, *SS*_*d*_ the self-sufficiency ratio for *d* is given by the following equation:

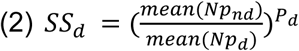

SS_*d*_ is thus centred around 1, increases if *d* has fewer partners than the other drivers, and the deviation from 1 is lessened when statistical power is low due to few observations for *d*. In order to avoid infinite ratios when the mean of *Np*_*d*_ is 0 due to the absence of any partner in all samples where d is mutated, *mean(NP*_*d*_) is substituted by (1 / (*length(Np*_*d*_) + 1)) in these cases. Only three such cases occurred, all in kidney clear cell carcinoma (KIRC) with ELF1, HDAC9 and SHMT1 mutations.

Epistatic interactions were defined as the ratio of observed to expected number of pairwise co-occurrences between driver alterations in a tumour type, weighted by the distribution of mutations across samples and the number of sample bearing either mutation. To estimate the number of expected number of occurrences of any two alterations A and B, we first calculate the individual probabilities that A and B are mutated in each sample of the cohort given their relative frequency (i.e. number of A or B mutations divided by total number of mutations) and the number of mutations in each sample, using hypergeometric probabilities (phyper R function). This takes into account that co-occurrences are more likely to be observed in hypermutated samples. The probability that A and B co-occur in each sample *s* is thus given by P_*s*_(A&B) = P_*s*_(A) × P_*s*_(B), assuming that A and B are independent. We proceeded to 10,000 simulations for each combination: a draw is performed for each sample *s* with a probability of success equal to *P*_*s*_(A&B), and we recorded the number of success over all samples for each simulation. The co-occurrence ratio of A and B *C*_*ab*_ is given by the actual number of samples in which A and B co-occur divided by the mean number of successes in the 10,000 simulations. If the observed number of co-occurrences was 0, it was set to either 0.5 if the mean of all draws was > 0.5, or to the mean of the draws otherwise, so that the ratio would be 1, indicating an interaction neither positive nor negative due to the lack of sufficient observations. As for self-sufficiency, we weighted this score so as to penalise the few outliers observed when two very infrequent genes co-occur potentially by chance. This was done by multiplying *C*_*ab*_ by the fraction of the cohort that presented a mutation either in A or B.

### Scoring Local Malignant Adaptation: models of parameter combination

As the prevalence and interplay of each parameter is unknown, we designed different models that correspond to different combinations of the three LMA parameters investigated. For simplicity, all models rely on summing the contribution of each driver alteration to the adaptation of each individual sample, given its alteration load and tumour-type-specific context. Parameters in a model are then assigned specific weights, which are optimised so as to minimise the variation of the overall score between samples in each tumour type.

We separated the 3 parameters into two types of LMA components: *intrinsic* (selective advantage, self-sufficiency) and *interactive* (epistatic interactions). In any given tumour type, the intrinsic component is driver-specific, while the interactive component is context-dependent and relies on which other driver alterations are also present in a sample. We designed 4 models based on the combination of 2 criteria: 1) whether selective advantage and self-sufficiency are separate or combined; 2) whether the epistatic score of a driver is given by the mean of all its interactions with the other drivers in the sample, or by the product of these interactions. This thus corresponds to 4 possible designs in total, which are labelled separate_mean, separate_prod, combined_mean, combined_prod (Supp. Table 1). The former 2 models therefore comprise the 3 parameters weighed individually then summed, while the latter 2 models only comprise 2 individually weighted parameters, with the intrinsic component being given for each driver *d* by *SSd* multiplied by *SA*d. Such a multiplicative relationship is more relevant than an additive one, as *SAd* (range: 0.3 − 208.5) has higher variability than *SSd* (range: 0.3 − 1.8).

### Weight inference

To assign weights to the components of each model, we used 37 possible empirical values ranging from 0.01 to 100, symmetrically mirrored around 1. The complete list is [0.010, 0.011, 0.012, 0.014, 0.017, 0.020, 0.025, 0.033, 0.05, 0.10, 0.11, 0.12, 0.14, 0.17, 0.20, 0.25, 0.33, 0.5, 1, 2, 3, 4, 5, 6, 7, 8, 9, 10, 20, 30, 40, 50, 60, 70, 80, 90, 100]. For all 4 models, we performed LMA score calculations using all possible combinations of values for each of the model’s parameters. For instance, using the combined_prod model with a 0.2 factor for the intrinsic component and the 3 factor for the interactive one, the LMA score for a sample *s* with *N* drivers is given by the following equation:

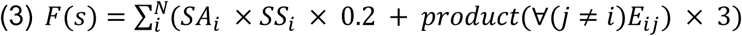

Where *E*_*i*j_ is the epistatic score for the interaction of drivers *i* and *j*. *SA*_*i*_, *SS*_*i*_ and *E*_*ij*_ depend on the context given by the tumour type of *s*. We calculated the standard deviation of the LMA scores from the related samples normalised by their median. We used an objective function aiming to minimise the standard deviation in normalised LMA while assessing all weight combinations, so as to match our initial assumption that most tumour-initiating clones in a given environment have a similar score. As different tumour types can involve different development mechanisms, the weights were optimised in tissue-specific fashion. The weight combination yielding the lowest standard deviation in normalised LMA was thus selected in each tumour type.

### Network representation

Network images were produced using the cytoscape software (Shannon et al., 2003).

## Results

### Cancer evolution is highly repeatable

In order to calculate the adaptation of a genetic clone to its local environment, we defined landscapes of Local Malignant Adaptation (LMA) specific to each of 9 cancer types, based on the occurrence of their most frequent driver alterations. To accurately reflect the genetic background of the clones that ultimately adapted to each environment, we only focused on clonal driver alterations (i.e. those present in all cells of a tumour). We investigated the repeatability of cancer evolution in each tumour type using a Jaccard Index based at the gene level, considering all sample-specific sets of clonal alterations as the genotypes of similarly adapted phenotypes (Bailey, Rodrigue, & Kassen, 2015; Yeaman, Gerstein, Hodgins, & Whitlock, 2018). Results were compared to randomised distributions of mutations that followed the same mutational load per patient (see Methods). As expected given the recurrence of driver mutations, our results highlight a high parallelism within each tumour type (Fig. 1A). On average, the similarity between samples from the real distribution of alterations were 1.4 to 5.1 times superior to the 95th percentile of those observed in our randomised controls (Fig. 1B). The glioblastoma (GBM) and lung squamous cell cancer (LUSC) sets displayed particularly high parallelism, with 96% of the indices being higher than the 95th percentile of the randomised control.

**Fig. 1.**
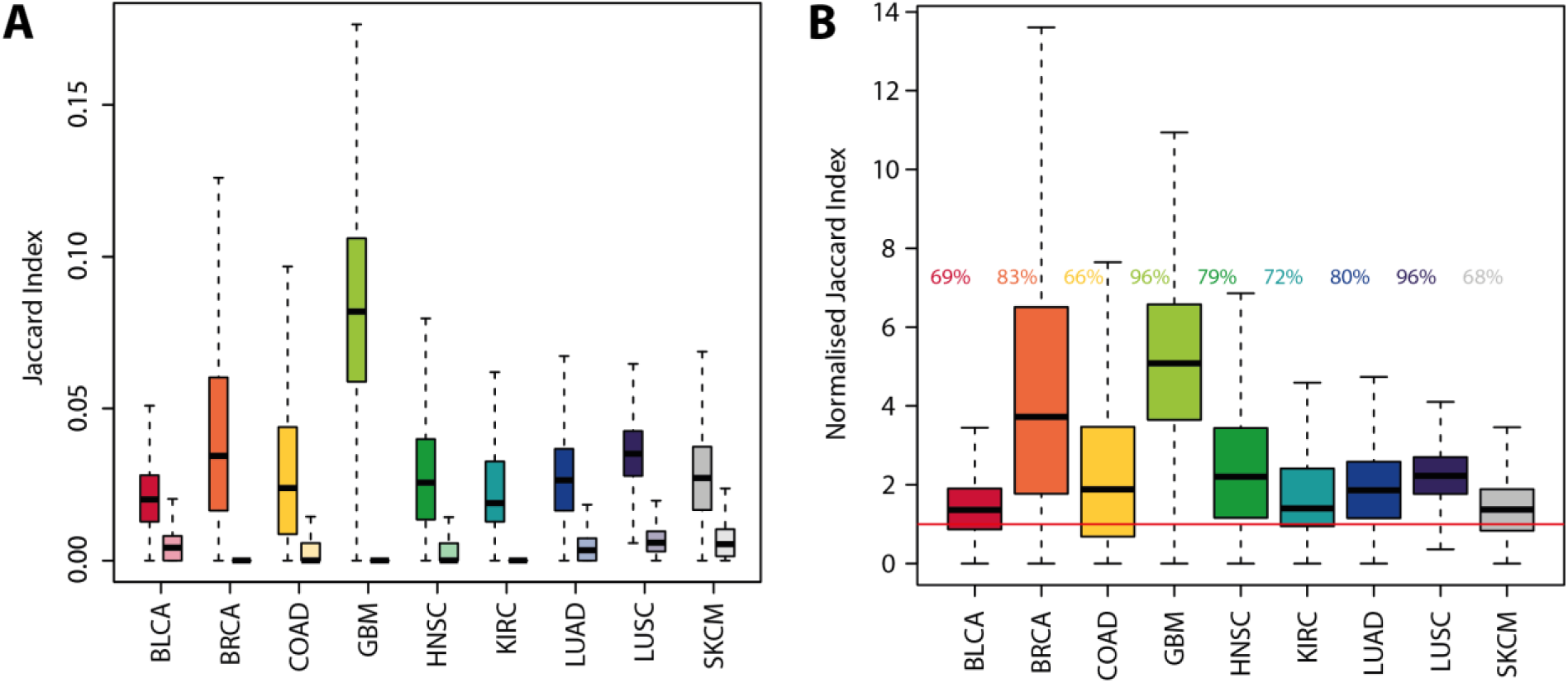
Repeatability of cancer evolution. A) Jaccard distances between the clonal genetic makeup of samples in the TCGA (left, vivid colours) and randomised controls (right, light colours) for each of the 9 tumour types. B) Jaccard distances in the TCGA data normalised via division by the 95^th^ percentile of each corresponding randomised data. Horizontal red line highlights a value of 1 (no difference). The percentage of observations exceeding the simulated 95^th^ percentile is reported left of each distribution. boxes represent the middle quartiles; black horizontal bars represent the median of each distribution; whiskers extend up to 1.5 times the interquartile range (box height) away from the box. Outliers (beyond the whiskers) are not displayed.

### Genetic landscapes of Local Malignant Adaptation: parameter definition

We identified 3 types of genetics-based factors that could quantify the adaptation of cells to a determined environment: selective advantage, driver self-sufficiency and epistatic interactions (Fig. 2A). Selective advantage has been the focus of numerous previous studies (Gonzalez-Perez & Lopez-Bigas, 2012; Lawrence et al., 2013; Martincorena et al., 2017; Zapata et al., 2018), and was defined in our case as the ratio of observed occurrence of each clonal alteration (driver or not) compared to its expected occurrence given its number of clonal observations, weighted by gene size (in base pairs). We compared our selection score to the corrected dN/dS measure obtained by Zapata et al. in a recent publication (Zapata et al., 2018). We find that both scores are moderately, yet significantly correlated (p < 0.001, Supp. Fig. 1). The observed variability can furthermore be explained by the fact that their analysis was based on pan-cancer data while our genetic landscapes are tumour-type specific, and the fact that we focus solely on clonal alterations.

**Fig. 2.**
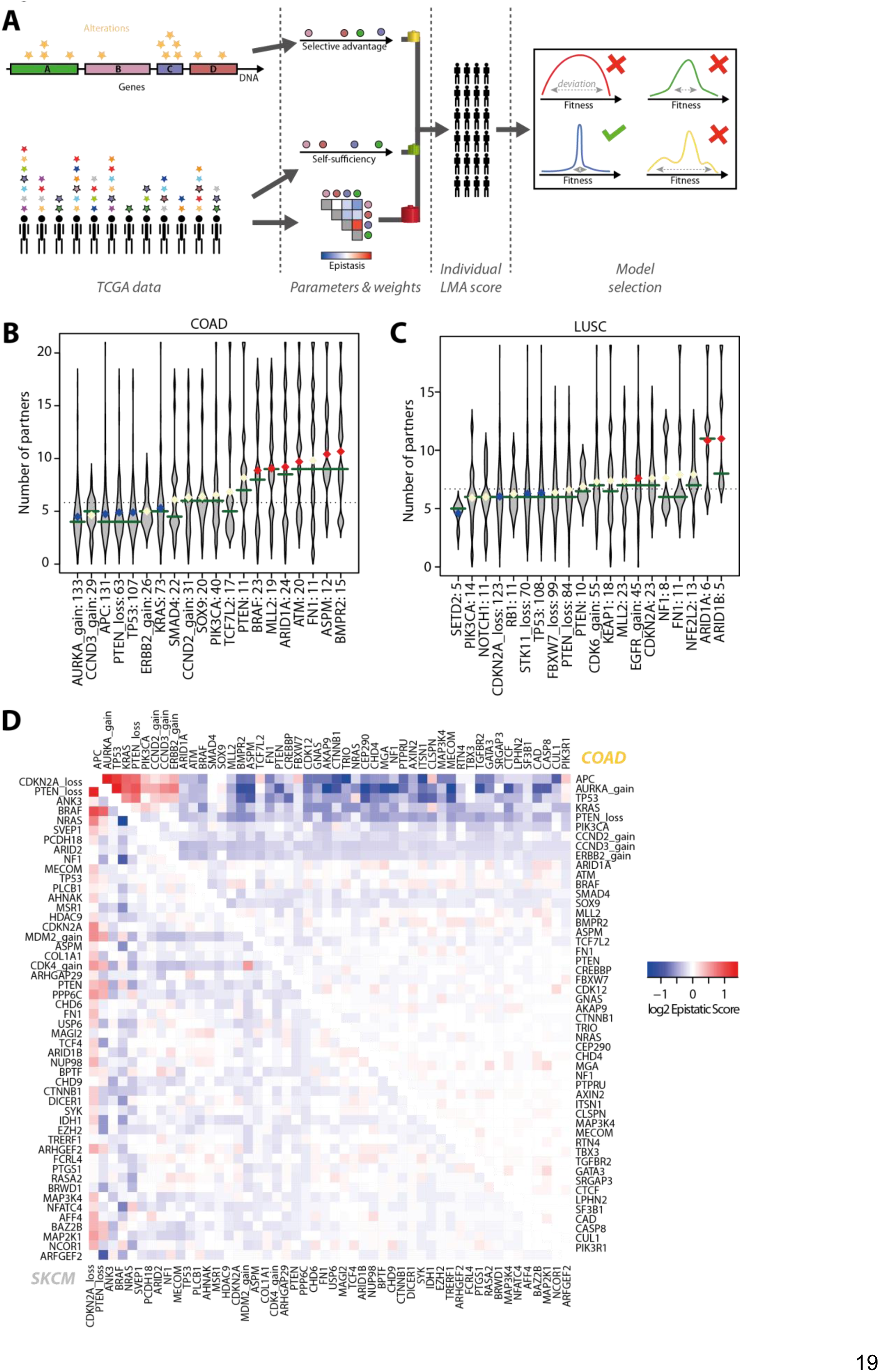
Evolutionary parameters of Local Malignant Adaptation. A) General scheme. Selective advantage is computed from the repartition of alterations per gene, self-sufficiency and epistatic interactions are calculated from the repartition of alterations per patient. The parameters are combined according to different models to calculate the LMA of all tumour-initiating clones in each cohort. The parameters corresponding to the lowest deviation in the overall LMA score across specimens are selected. B) Number of additional drivers observed in samples harbouring the 20 most frequent alterations in COAD (colorectal adenocarcinoma) and C) LUSC (lung squamous cell carcinoma). Horizontal green bars represent the median; diamonds represent the mean. Blue means significantly fewer partners than expected, red means significantly more. Dotted line indicates the overall mean. D) Epistatic interactions between all retained drivers in COAD (top right) and SKCM (skin melanoma, bottom left). Blue indicates negative interaction due to co-occurrences rarer than expected, red indicates positive interactions (higher co-occurrence than expected).

Driver self-sufficiency is however a novel measure, reflecting how many additional drivers are needed on average to induce malignant development. Differences could be observed across tumour types, with for instance a high discrepancy in the number of additional drivers in colorectal adenocarcinomas and a relatively homogeneous distribution in lung squamous cancers (Fig. 2B-C, Supp. Fig. 2).

Both selective advantage and self-sufficiency measures are specific to single driver alterations in a given tumour type. We found they were significantly correlated, yet with a very high variability (R^2^=0.04, p<0.001, Supp. Fig. 2). This indicates that they can provide distinct information on the impact of each alteration and that mutations highly selected for may still require numerous other alterations in order to induce full cancerous transformation. We defined epistatic interactions as the ratio of co-occurrence between two genes given their respective frequencies and the number of mutations per sample in a cohort, using hypergeometric probabilities (see Methods). The most negative interaction was the one found between known antagonists BRAF and NRAS in melanoma (Curtin et al., 2005), while the most positive interactions included those between AURKA gain, APC and TP53 mutations in colorectal cancer (Fig. 2E). Across cancers, CNAs tended to frequently co-occur with each other and TP53 mutations (Supp. Fig. 4), in agreement with the role of TP53 in promoting genome instability, which then accelerates CNA acquisition (Martinez et al., 2016; Sansregret, Vanhaesebroeck, & Swanton, 2018).

### Model selection

In this study, we aim to calculate the adaptation of genetic clones that reached invasive potential in each tumour type. We decide to model LMA as the sum of the contribution of all clonal drivers in a sample using different models, corresponding to distinct combinations of the 3 parameters previously calculated. The selective advantage and self-sufficiency parameters correspond to the intrinsic component of LMA, as they solely depend on individual driver properties. Epistatic interactions define the interactive component of LMA, depending on interaction between all drivers present in a sample. We used four models based on these two criteria: either combining (“combined”) or separating (“separate”) selective advantage and self-sufficiency; and either quantifying the epistatic interactions as the mean of all interactions between a driver and its partners (“mean”) or the product of these interactions (“product”). We then established the optimal weights for all components of each model that minimised the variation in LMA across all samples on a tumour type basis (see Methods). This objective function aims at producing data matching our assumption that the initiating clones in different tumours of the same type are all similarly well adapted in this landscape.

We evaluated the relevance of these models by calculating the prevalence of each component in the LMA score of all samples on a per set basis, as well as the correlation between LMA and the number of drivers in each sample (Fig. 3). Both models separating the intrinsic component of LMA were found to be optimal by down-weighing selection strength and the resulting scores were often dominated by a single component, often being self-sufficiency (Fig. 3A-B, Supp. Fig. 5). In addition, the separate_mean model diminished the contribution of epistatic interactions, while they appeared as major contributors in several tumour types under the separate_prod model. A similar observation was made for the combined models, in which the epistatic interactions were down-weighted in the combined_mean model while this was not the case in the combined_prod model (Fig. 3C-D). Under all models and in most tumour types, the resulting LMA score of a sample was highly correlated to its number of drivers (all p<0.001, Fig. 3E). The combined_prod model was however the model in which LMA and number of drivers were the least correlated (p<0.01 against all other models, paired Wilcoxon test). We therefore elected the combined_prod model as the most appropriate of the 4 models to calculate LMA, as it was less dominated by a single component and was less likely to measure adaptation as a static process of stacking up driver alterations.

**Fig. 3.**
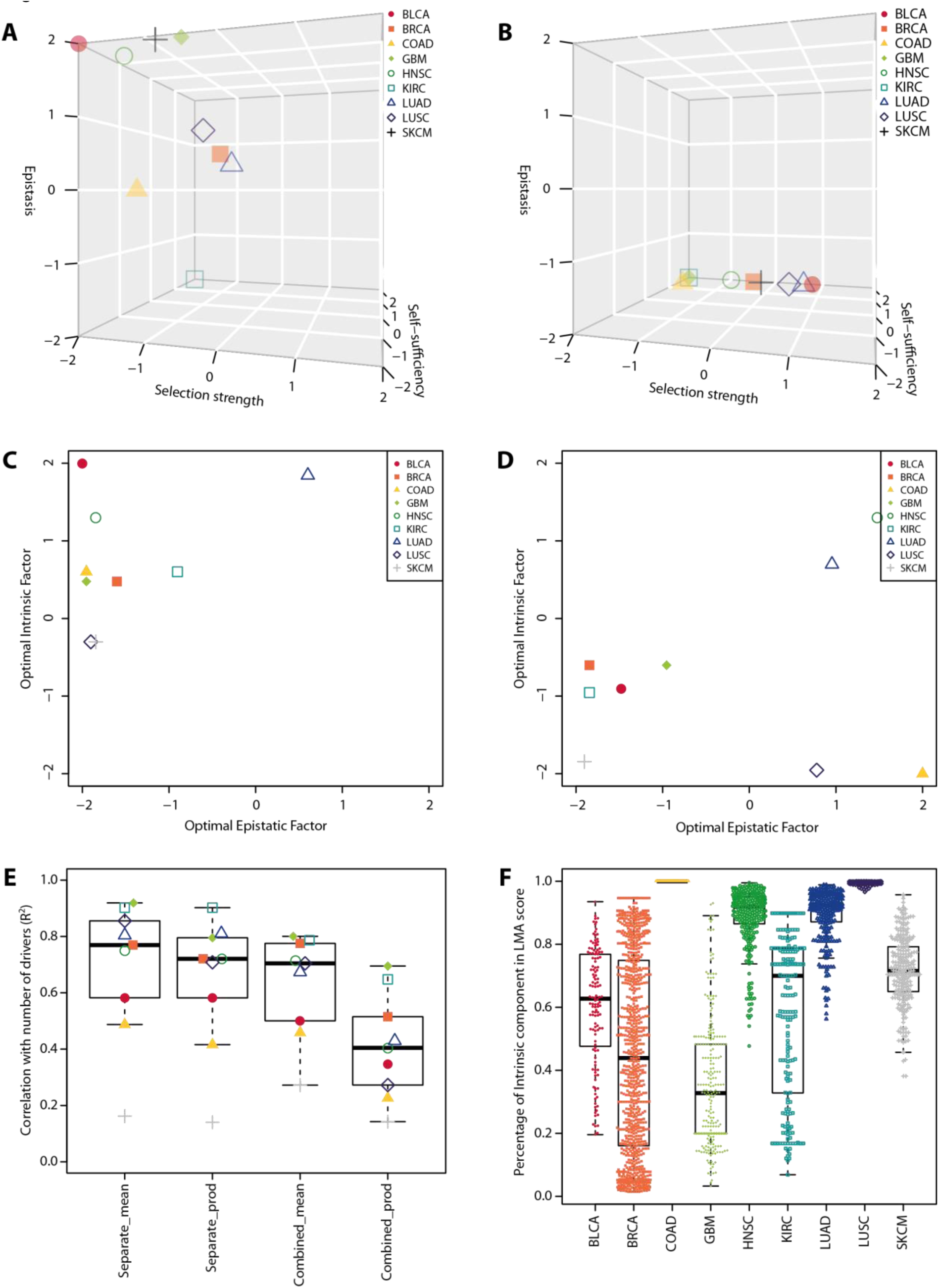
Model selection. A-B) Optimal weights found for selective advantage, self-sufficiency and epistatic interactions in the “separate” models (log10 scale). A) separate_mean; B) separate_prod. C-D) Optimal weights found for the intrinsic (selective advantage x self-sufficiency) and interactive (epistatic interactions) components of the combined models (log10 scale). C) combined_mean; D) combined_prod. E) Distribution of R^2^ values for the correlation between Local Malignant Adaptation and number of drivers in all 9 tumour types according to all 4 models. F) Share of the LMA score of each tumour specimen corresponding to the intrinsic component using the combined_prod model in all tumour types.

### Differences among and within cancer types

We measured the percentage of the LMA scores accountable to the intrinsic component for each sample of each tumour type (Fig. 3F). Our results suggest our assumption that all malignancies are equivalently adapted to their local environment could rely on different evolutionary dynamics in different tissue-specific contexts. Colorectal, head & neck and lung (adenocarcinomas and squamous cell) cancers were defined by a prevalence of the intrinsic component in the scores of their specific samples, while the interactive component prevailed in glioblastoma. Interestingly, we observed a high variability in the share of each LMA score component of all samples in many tumour types, particularly in breast cancer. These observations reflect the extensive heterogeneity recurrently observed both inter- and intra-tumour at the genetic and phenotypic levels, which can be mirrored by our occurrence-based framework to calculate local malignant adaptation.

### Pre-malignant lesions are specifically adapted to the local landscape

We analysed two published cohorts of pre-malignant lesions linked to melanomas and colorectal carcinomas (CRC), to understand whether these precursors differed from full-blown tumours in our measure of adaptation. We first analysed 30 malignant and 23 benign skin lesions from (Shain et al., 2015), including 18 precursor/melanoma pairs. We observed that the LMA score of the melanomas was consistently and significantly higher than the one of their paired benign precursor (p=0.001 paired Wilcoxon test, p=0.01 unpaired, Fig. 4A). We additionally analysed 9 colorectal adenomas and 11 carcinomas from (*Cross et al, under review*). Multiple samples were available for all cases (2-6 per adenoma, 4-13 per carcinoma, see Methods). The LMA of carcinomas was higher than the one of adenomas, although not significantly, possibly due to the limited sample size and absence of benign/malign pairing (p=0.37, Wilcoxon test, Fig. 4B). A similar trend was observed with significance when investigating individual biopsies rather than whole lesions, although the redundancy and uneven number of samples across cases may bias this observation (p=0.018, Wilcoxon test, Supp. Fig. 6).

**Fig. 4.**
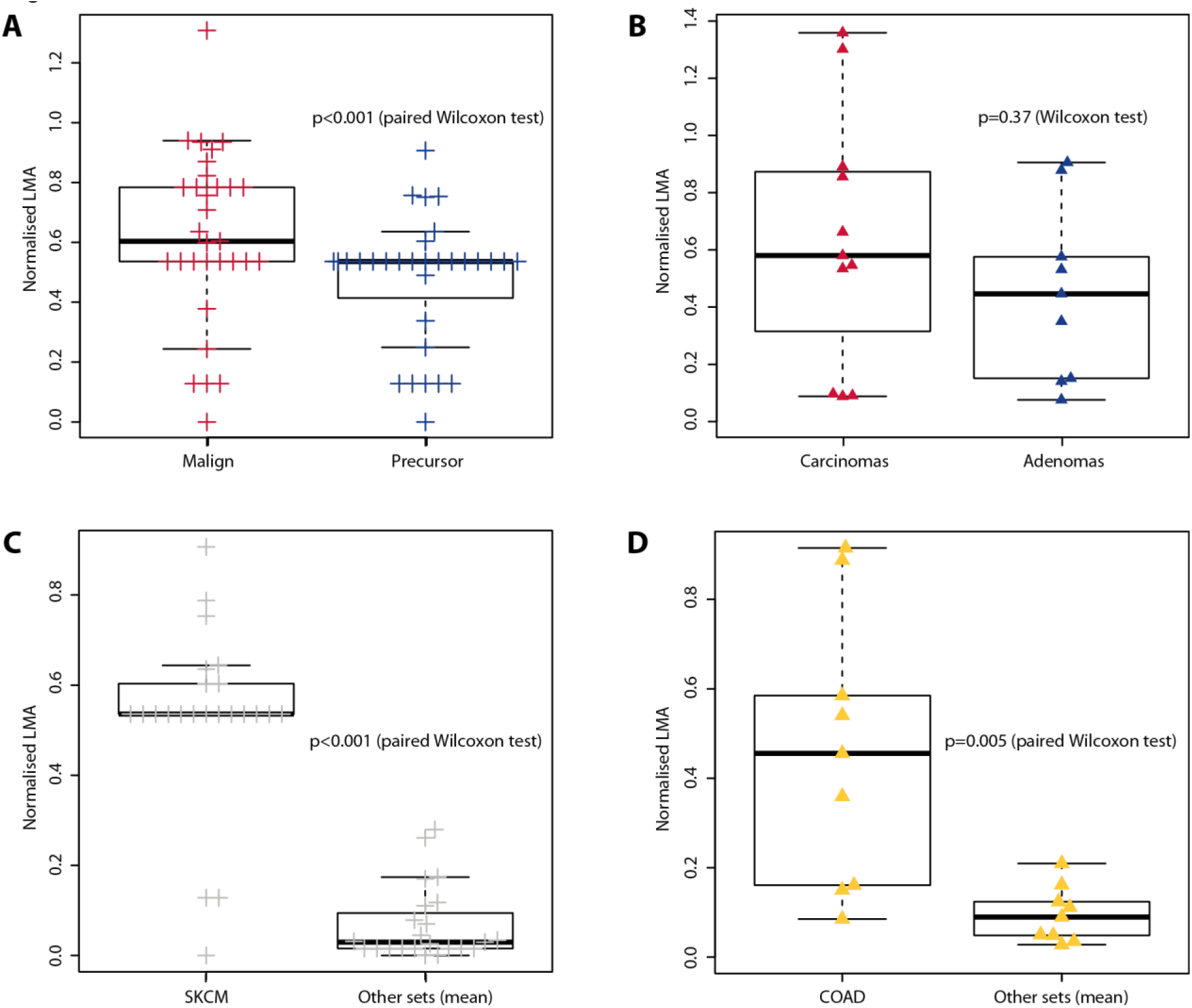
Pre-malignant lesions. A) Normalised LMA scores of paired melanomas (left) and their precursor lesions (right). B) Normalised LMA scores of unpaired colorectal carcinomas (left) and adenomas (right). C) Normalised LMA scores of nevi in the SKCM landscape (left) and in the 8 other landscapes on average (right). D) Normalised LMA scores of adenomas in the COAD landscape (left) and in the 8 other landscapes on average (right).

When we calculated the LMA scores of the 23 nevi and 9 adenomas in other tumour-type-specific landscapes, they appeared more adapted respectively to the melanoma and CRC landscapes than to the other landscapes on average (p<0.001 and p=0.005, paired Wilcoxon test, Fig. 4C-D, Supp. Fig. 7). Our model is thus able to detect that pre-malignant lesions are specifically adapted to their local environment, further suggesting that additional evolutionary steps are still necessary to acquire locally invasive capacity.

### Metastasis does not rely on genetic adaptation to the distant site’s landscape

We then applied our methods to metastatic samples, in order to shed light on whether the genetic basis of adaptation to an environment was as relevant for its metastatic colonisation as for local invasion. We used the MET500 dataset of 500 metastatic samples (Robinson et al., 2017) and used sample annotation to identify 170 samples for which we had a primary site LMA landscape (i.e. one of the 9 tumour types analysed), 82 samples with an available metastatic site landscape, and 35 for which the landscape of both metastatic and primary sites was known. Manual curation was employed to attribute a relevant tumour type to each sample (Supp. Tables 2 and 3). As we had 2 different potential landscapes corresponding to lung metastases (LUAD, adenocarcinoma, and LUSC, squamous cell cancer), all calculations were replicated by taking either type as default landscape for lung metastases. For the 35 samples with existing landscapes for the primary and metastatic sites, we observed that LMA scores were consistently higher in the primary site than in the metastatic one (p<0.001, paired Wilcoxon test, Fig. 5A and Supp. Fig. 8). This suggests that metastatic clones are more genetically adapted to the environment in which they originated than to the one they colonised. As for pre-malignant lesions, we analysed the LMA scores of each sample in all the other tumour types. LMA in the landscape of the primary site was significantly higher than the average LMA in the other 8 landscapes (p<0.001, paired Wilcoxon test, Fig. 4B, Supp. Fig. 9), while we did not observe any difference when considering the adaptation in the metastatic site’s landscape (Fig. 5C, Supp. Fig. 10). This suggests that these genetic clones were specifically adapted to their environment of origin. However, their genetic makeup did not provide them with the potential to specifically adapt to their metastatic site as well as local primary tumours.

**Fig. 5.**
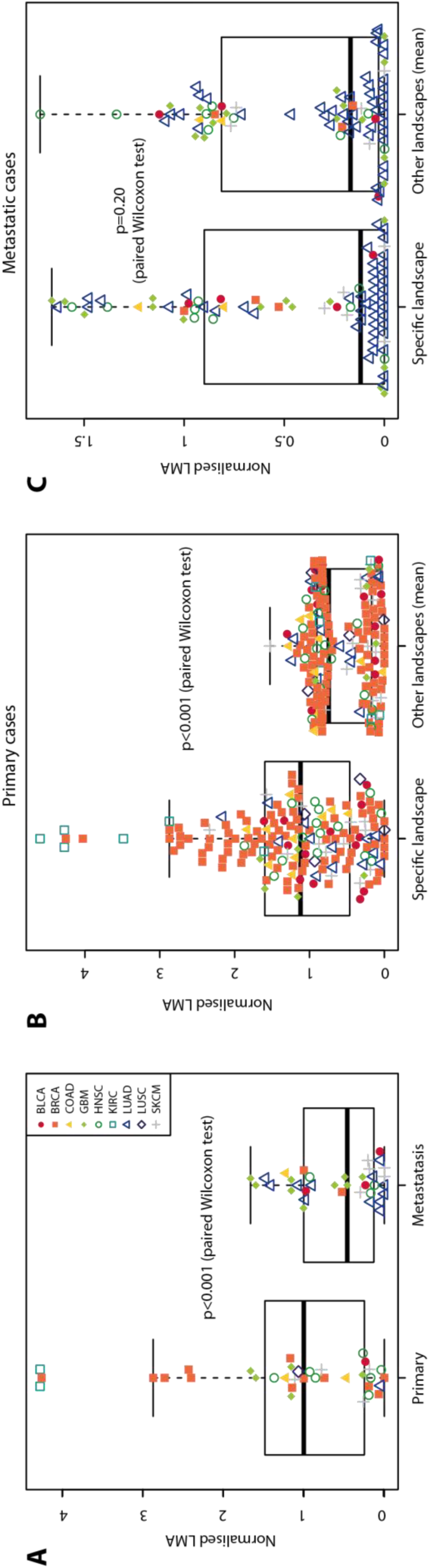
Metastatic lesions. A) Normalised LMA scores of 35 lesions with landscapes existing for both primary and metastatic site. Left, LMA in the primary site’s landscape; right, LMA in the metastatic site’s landscape. B) Normalised LMA scores of 170 lesions with a primary site landscape in their specific landscape (left), or in the other 8 landscapes on average (right). C) Normalised LMA scores of 82 lesions with a metastatic site landscape in their specific landscape (left), or in the other 8 landscapes on average (right).

### TCGA landscapes as graphs

Our results demonstrate that our LMA model can quantify genetic adaptation to specific somatic evolutionary contexts. Our method can further be combined to a network approach to understand adaptation dynamics as individual driver alterations accrue in a clone over time. Fitness landscapes can be represented by graphs (Diaz-Uriarte & Wren, 2018; Palmer et al., 2015) in which nodes are unique combinations of drivers, each with a distinct fitness. We reproduced such and architecture with our LMA scores, where edges connect nodes as additional drivers are added on top of previous combinations (Fig. 6A). This network architecture can represent all evolutionary trajectories as successive acquisitions of any given number of driver alterations. As such a combinatorial approach is computationally heavy, we restricted our analysis to the 15 most common alterations in each tumour type. This corresponds to a maximum of 32,767 unique combinations of driver alterations per tumour type.

**Fig. 6.**
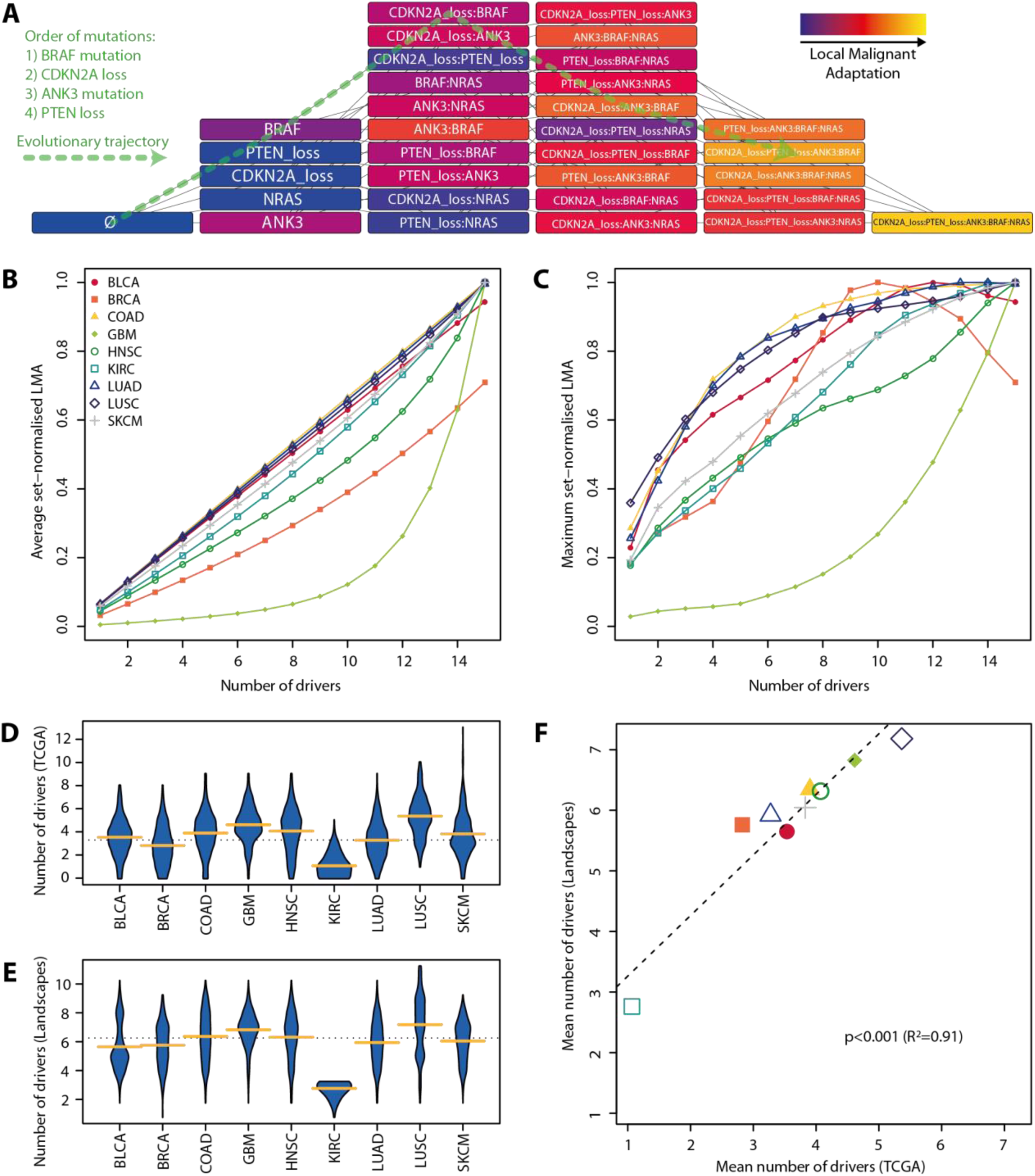
Graphs of Local Malignant Adaptation. A) Example graph with the 5 most common driver alterations in colorectal cancer (COAD). Nodes are combinations of alterations, each connected to all possible previous and posterior combinations of step-by-step driver acquisition. LMA is increasingly coloured in blue to red to yellow. An example of evolutionary trajectory in which a clone subsequently acquires 4 mutations is highlighted by a green dashed line throughout the network. B) Average and C) maximum node LMA score per number of driver alterations in all tumour types. Landscapes were limited to the 15 most prominent drivers of each tumour type. D) Number of clonal drivers per TCGA patient and E) number of drivers in all evolutionary trajectories allowing to reach a minimal LMA score equal to the median TCGA score of the same tumour type. Yellow bars indicate the mean of each tumour type, dotted line indicates the overall mean. F) Correlation between the mean number of clonal drivers actually observed in TCGA patients and the mean number of drivers required to reach at least the median TCGA score in the corresponding landscape.

We took advantage of this framework to investigate how malignant adaptation to the local environment evolves in the 9 investigated TCGA tumour types, by following how genetic clones progress through the graph by acquiring new alterations. We observed differences in LMA dynamics depending on the properties of each tumour type. We see that in many cases LMA increases linearly on average with each novel driver, as can be expected from our approach based on summing the contextualised contributions to adaptation of each driver in a clone. However, tumour types in which our model predicted a high prevalence of epistatic interactions (GBM, BRCA) display a strong deviation from linearity (Fig. 6B). This is also reflected in how fast the maximum LMA score in the network is reached, with some tumour types displaying a log-like distribution with decreasing improvement with each additional alteration, while the maximum LMA of GBM increases exponentially (Fig. 6C). Interestingly, the maximum LMA for BLCA, BRCA and COAD is reached early and decreases after respectively 10, 12 and 13 alterations, as would be expected under diminishing returns epistasis (Chou, Chiu, Delaney, Segrè, & Marx, 2011). This highlights that our model can produce non-linear relationships between LMA and the number of drivers, depending on the nature of epistatic interactions among driver alterations.

Finally, we investigated if these network representations could represent a basis on which to predict the evolutionary trajectories potentially leading to cancer. In all tumour types, we recorded all trajectories that reached a LMA score superior or equal to the median score observed in the corresponding TCGA cohort. The first node meeting such a criterion in a trajectory was considered final and its offspring nodes were not investigated. These trajectories thus represent all combinations of genes that likely lead to sufficient genetic adaptation for tumorigenesis. The number of drivers in these combinations was superior to the one observed in the actual sample, which was expected given that their LMA score had to be equal or higher than the dataset’s median (Fig. 6D-E). The mean number of required alterations is however strongly correlated to the mean number of clonal drivers observed in each tumour type (R^2^=0.91, p < 0.001, Fig. 6F). This suggest that networks of contingency-based adaptation metrics can thus provide a framework to both represent and study pre-cancerous progression under a novel angle, while recapitulating the specificities of different tumour types. Their use can furthermore identify system-level properties of the evolutionary context that funnels malignant somatic evolution in different organs and environments.

## Discussion

Here we developed a methodology to investigate the progression of normal cells towards oncogenesis, by way of measuring the adaptation of genetic clones to a given environment. Our method relies on the contingency and repeatability of cancer development, observed when multiple same-type cancers occur with distinct evolutionary trajectories in different human individuals. We built and optimised simple models to quantify Local Malignant Adaptation based on the presence of recurrent genetic alterations in 9 cohorts corresponding to 9 tumour types. This approach is similar to fitness landscapes that aim to map the space in which phenotypes evolve and adapt in evolutionary biology. We then applied our method to independent datasets of pre-malignant and metastatic lesions to analyse the impact of genetics in the adaptation to and the colonisation of different environments.

Our exploratory model aimed at simplicity and still suffers from several limitations. First of all, our model is solely based on genetics and ignores the potential influence of a cell’s phenotypic state on its response to specific genetic insults, which can impact tumorigenesis (Morel et al., 2017; Puisieux, Pommier, Morel, & Lavial, 2018) or lead to non-genetic selection (Shaffer et al., 2017). It also ignores the contribution of epigenetic alterations as potential drivers, the inclusion of which will require a more general understanding on the recurrent epigenetic alterations functionally linked to tumour formation (Timp & Feinberg, 2013). There furthermore exists no method to estimate the clonality of CNAs, thus hampering our accuracy in including only the truly clonal ones in LMA calculations. Despite the fact that our model includes epistatic relationships, it is unable to estimate the ordering of alterations, which can heavily influence evolutionary trajectories (Ortmann et al., 2015). Aside from genetic alterations, our model does not include the interactive adaptation relationship between a pre-invasive cell and its environment, the interplay between both being very likely to modify selective pressures as potential tumour-initiating cells develop (Bissell & Radisky, 2001; Rozhok, Salstrom, & DeGregori, 2016; Scott & Marusyk, 2017). Finally, our work focuses on a single clone being responsible for initial local invasion, while it is possible that this process can involve multiple clones (Casasent et al., 2018). Such polyclonal invasion would however likely involve a recent common ancestor. Addressing these drawbacks in the future will allow to better stochastically model malignant progression.

Despite these limitations, our model fits the hypothesis that carcinogenesis is likely a stepwise process. Our analyses highlighted that pre-malignant lesions were specifically adapted to their environment, yet insufficiently to promote local malignant invasion. This thus suggests that cancer arises when benign lesions acquire further driver alterations, at least in melanomas and colorectal carcinomas. In addition, the analysis of metastatic lesions suggested that genetics contributed to the formation of the primary tumours but were not a defining factor in the adaptation to the metastatic site. Metastasis to a specific site requires convergent evolution for clones from divergent backgrounds to adapt to the new environment (Cunningham, Brown, Vincent, & Gatenby, 2015). This adaptation may however not depend on novel genetic alterations, potentially relying on cellular plasticity (Varga & Greten, 2017), or may involve completely different genetic determinants than ab-initio oncogenesis in the same site.

Finally, we propose that this methodology could be combined with network approaches to model oncogenesis as a stochastic process and investigate systemic differences between tumour types. We report that our approach reflects the specificity of tumorigenesis in the different evolutionary contexts dictated by the tissue of origin. Future improvements could help the accurate reconstruction of tumorigenesis in-silico, allowing to computationally investigate new opportunities for early detection and prognostication. A recent method based on phylogenetics and machine-learning also detected patterns in the repeatability of cancer evolution, which can help classify patients on their prior (and likely future) evolutionary trajectories (Caravagna et al., 2018). By extension, reconstructing fitness/adaptation landscapes of already invasive tumours based on different therapeutic options can help tailor ad-hoc therapeutic regimens: based on the fitness and frequency of all (sub)clones composing each tumour, these landscapes can allow to optimise sequential drug schedules so as to dynamically reduce overall fitness (Nichol et al., 2017). Occurrence-driven definition of epistatic interactions requires large numbers of observations but can be completed by animal models in which driver combinations can be induced and followed over time (Rogers et al., 2018). The reconstruction of therapy-specific human fitness landscapes thus critically requires efforts to generate large, centralised public datasets, with accurate clinical annotation for each treatment type, ideally with samples before and after treatment.

## Author contributions

PM designed the experiments. NT and PM analysed the data. CML and SS organised the collaboration. CML, AP, SS and PM supervised the work. PM wrote the article. All authors reviewed the article.

## Acknowledgements

The results published here are in part based upon data generated by the TCGA Research Network: http://cancergenome.nih.gov/. International mobility and collaboration were facilitated by the International Alliance Research Internship (IARI, part of the Programmes d’Investissement d’Avenir) programme between the universities of Tokyo (UTokyo) and Lyon (UdL). The authors wish to thank William CH Cross and Trevor A Graham for their help with the colorectal data; as well as Benjamin Roche and Robert J Noble for valuable suggestions on the manuscript.

## Data archiving statement

The code and data used for this analysis are available on github: https://github.com/pierremartinez/ConFitLand.

